# Auditory Hypersensitivity and Processing Deficits in a Rat Model of Fragile X Syndrome

**DOI:** 10.1101/2021.09.25.461569

**Authors:** Benjamin D. Auerbach, Senthilvelan Manohar, Kelly Radziwon, Richard Salvi

## Abstract

Fragile X (FX) syndrome is one of the leading inherited causes of autism spectrum disorder (ASD). A majority of FX and ASD patients exhibit sensory hypersensitivity, including auditory hypersensitivity or hyperacusis, a condition in which everyday sounds are perceived as much louder than normal. Auditory processing deficits in FX and ASD also afford the opportunity to develop objective and quantifiable outcome measures that are likely to translate between humans and animal models due to the well-conserved nature of the auditory system and well-developed behavioral read-outs of sound perception. Therefore, in this study we characterized auditory hypersensitivity in a *Fmr1* knockout (KO) transgenic rat model of FX using an operant conditioning task to assess sound detection thresholds and suprathreshold auditory reaction time-intensity (RT-I) functions, a reliable psychoacoustic measure of loudness growth, at a variety of stimulus frequencies, bandwidths and durations. Male *Fmr1* KO and littermate WT rats both learned the task at the same rate and exhibited normal hearing thresholds. However, *Fmr1* KO rats had faster auditory RTs over a broad range of intensities and steeper RT-I slopes than WT controls, perceptual evidence of excessive loudness growth in *Fmr1* KO rats. Furthermore, we found that *Fmr1* KO animals exhibited abnormal perceptual integration of sound duration and bandwidth, with diminished temporal but enhanced spectral integration of sound intensity. Because temporal and spectral integration of sound stimuli were altered in opposite directions in *Fmr1* KO rats, this suggests that abnormal RTs in these animals are evidence of aberrant auditory processing rather than generalized hyperactivity or altered motor responses. Together, these results are indicative of fundamental changes to low-level auditory processing in *Fmr1* KO animals. Finally, we demonstrated that antagonism of metabotropic glutamate receptor 5 (mGlu5) selectively and dose-dependently restored normal loudness growth in *Fmr1* KO rats, suggesting a pharmacologic approach for alleviating sensory hypersensitivity associated with FX. This study leverages the tractable nature of the auditory system and the unique behavioral advantages of rats to provide important insights into the nature of a centrally important yet understudied aspect of FX and ASD.

## 1. Introduction

Human genetic studies have greatly increased our understanding of the gene mutations associated with the increased prevalence of autism spectrum disorders (ASD) (de la Torre-Ubieta et al., 2016; Doan et al., 2019). The results have facilitated the development of genetically validated animal models, which have been instrumental in the identification of the cellular and molecular disturbances linked with ASD (Moy and Nadler, 2008; Schroeder et al., 2017). Connecting molecular pathologies to behavioral phenotypes of ASD, however, remains a significant challenge that has impeded the development of ASD therapies (Berry-Kravis et al., 2018; Vorstman et al., 2017). A case in point is Fragile X syndrome (FX), the leading inherited cause of ASD (Hagerman et al., 2017). FX is caused by CGG expansions around the FMR1 gene, leading to its transcriptional silencing and subsequent loss of its protein product FMRP (Verkerk et al., 1991). The known genetics of FX and the evolutionarily conserved nature of FMRP have allowed for the development of well-validated animal models of FX that have provided important insights into its pathophysiological mechanisms (Berry-Kravis, 2014; Krueger and Bear, 2011). For instance, animal studies have demonstrated that dysregulated metabotropic glutamate receptor 5 (mGlu5) signaling is a core component of FX pathophysiology and mGlu5 inhibitors have been successful at ameliorating many cellular, synaptic, and behavioral phenotypes in FX models (Bhakar et al., 2012; Michalon et al., 2012; Pop et al., 2014). Despite this preclinical success, clinical trials targeting molecular disturbances in FX have been largely disappointing to date (Anagnostou, 2018; Berry-Kravis et al., 2018). Although many factors contribute to the challenges of clinical translation, one of most important gaps identified in pre-clinical animal studies is a lack of robust, clinically relevant behavioral phenotypes in animal models (Erickson et al., 2017). To address this gap, this study sought to develop a quantitative and disease-relevant behavioral read-out that could serve as a clinically translatable platform for screening potential therapies in FX models.

Sensory hypersensitivity and hyperreactivity are defining features of FX and ASD (Sinclair et al., 2017). One of the most common and debilitating sensory disturbances in FX and ASD is hyperacusis, an auditory hypersensitivity disorder in which moderate intensity sounds are perceived as unbearably loud (Danesh et al., 2015; Gomes et al., 2008; McCullagh et al., 2020; Rotschafer and Razak, 2014; Williams et al., 2021b). Loudness hyperacusis is not only an important clinical problem in FX and ASD, but may also provide a behavioral framework for the development of objective and quantifiable outcome measures that are likely to translate between humans and animal models due to the well-conserved nature of the auditory system and well-developed behavioral read-outs of sound perception. Operant sound detection tasks, where animals are trained to generate a behavioral response to specific stimuli, allow for detailed assessment of auditory detection speed and accuracy across a range of stimulus parameters using an experimental design that can be translated to human studies. Importantly, many of these psychoacoustic measures are also quantitative correlates for perceptual attributes of a stimulus. For instance, human and animal psychophysical studies have both shown that auditory reaction time (RT), the time it takes for a subject to respond to an acoustic stimulus, is inversely correlated with sound intensity. Reaction time-intensity (RT-I) functions collected in humans have been used to construct equal loudness contours that are well correlated with those obtained with subjective loudness scaling procedures (Marshall and Brandt, 1980; Melara and Marks, 1990; Seitz and Rakerd, 1997). RT-I functions collected in animals are predictably modulated by several acoustic parameters (frequency, duration, bandwidth) known to influence loudness judgements in humans (Green, 1975; May et al., 2009; Radziwon and Salvi, 2020; Stebbins, 1966). Thus, RT-I functions are an objective psychophysical read-out of loudness perception that is maintained across species. Indeed, RT-I functions have been used in several different animal models to quantify normal loudness growth (Stebbins, 1966), abnormal loudness growth due to cochlear hearing loss (Moody, 1973), and loudness hyperacusis resulting from salicylate ototoxicity or prolonged noise exposure (Auerbach et al., 2019; Radziwon et al., 2019; Radziwon et al., 2017).

Because auditory RT measures obey all the psychophysical rules of loudness perception with respect to intensity, frequency, stimulus bandwidth and duration, we used RT-I functions to carry out a comprehensive assessment of loudness perception in a transgenic rat model of FX containing a 122 bp deletion in exon 8 of the *Fmr1* gene (*Fmr1* KO rat) (Hamilton et al., 2014). This model of FX recapitulates core cellular pathophysiology of the disorder, including altered mGlu5 function (Till et al., 2015), but takes advantage of the highly trainable nature of rats to allow for in depth behavioral characterization not afforded by other model systems (Golden et al., 2019). Using RT-I measures, we found that male *Fmr1* KO rats exhibit increased loudness perception and disrupted spectral and temporal integration of loudness compared to WT controls, indicative of heightened sound sensitivity and disrupted auditory processing. Finally, we determined that normal RTs could be restored in *Fmr1* KO rats by inhibiting mGlu5 activity, demonstrating that this behavioral phenotype is related to a core molecular pathology of FX. Together, these results indicate that auditory RT differences may be a novel behavioral phenotype in FX that is directly related to core sensory disturbances in the disorder and can be used for preclinical screening of treatments.

## 2. Methods

### 2.1 Subjects

Adult (>2 month old) male *Fmr1*^*tm1sage*^ KO rats on an outbred Sprague-Dawley background (TGRS5390HTM4 FMR1 −/Y; SAGE Labs Inc., St. Louis, MO) and littermate wild-type (WT) controls were used for these studies. Male rats were used because FX occurs more frequently and in greater severity in males due to the X-linked nature of the disorder (Reiss and Hall, 2007). Nine *Fmr1* KO rats and nine WT littermates were used were used as subjects in most studies, except as noted. Rats were housed in pairs and maintained on a 12 h day/12 night cycle. The rats used in the experiments had free access to food and water except during operant conditioning, when rats were food restricted and kept at approximately 90% of their free-feeding weight. Rats in the operant conditioning studies were tested approximately 1 hr per day, 6-7 days per week. All experiments were approved by the University at Buffalo Institutional Animal Care and Use Committee (HER05080Y) in accordance with NIH guidelines.

### 2.2 Breeding and Genotyping

WT male rats (Charles River) were bred to heterozygous female *Fmr1* KO rats to generate male WT and *Fmr1* KO offspring used in these studies. Offspring were screened for a 122-base pair (bp) deletion in the *Fmr1* gene sequence using published procedures (Hamilton et al., 2014). Using a commercial kit (QIAGEN DNeasy isolation kit #69506), DNA was isolated from a tissue punch taken from the external ear. PCR was performed with a commercial kit (Sigma JumpStart™ Taq ReadyMix™, P2893) and 1-μL of purified DNA. The PCR amplification steps were: first cycle, 5 min at 95 °C; 35 cycles of 30-s each at 95 °C; 30-s at 60 °C; 40-s at 68 °C and a final cycle of 5-min at 68 °C. Primers used for amplification were S1 (5’ TGGCATAGACCTTCAGTAGCC 3’) and S2 (5’ TATTTGCTTCTCTGAGGGGG 3’). Primers were purchased from ThermoFisher Scientific. Amplified fragments were resolved in 2% agarose gel. The expected amplicon sizes on the gels were 400-bp for WT rats (+/+ or +/y), two products of 400-bp and 278-bp for heterozygotes (+/−) and 278-bp for homozygotes (−/− or −/y).

### 2.3 Operant psychophysical procedures

Rats were trained on a Go/No-go operant conditioning paradigm to detect sound bursts (5 ms rise/fall time, cosine gated) of varying intensity, frequency, duration, and bandwidth as described in our recent papers (Auerbach et al., 2019; Radziwon et al., 2019; Radziwon et al., 2017; Radziwon and Salvi, 2020). A rat started a trial by placing its nose in a nose-poke hole, which initiated a variable wait interval ranging from 1 to 4 s. The rat had to maintain its position in the nose-poke hole until it detected a sound or the trial was aborted (Fig. 1A). If the rat detected the signal and removed its nose from the nose-poke hole (Go condition) within a 2-s response interval, a food reward (45 mg dustless rodent pellets, Bio-Serv) was delivered and the response was scored as a HIT. A MISS was recorded if the rat failed to remove its nose from the nose-poke within the 2-s response interval. Approximately 30% of the trials were catch trials during which no stimulus was presented (No-go condition). If the rat kept its nose in the nose-poke during a catch trial, a correct rejection (CR) was recorded; no reinforcement was given for a CR, but another trial could be initiated immediately. If the rat removed its nose during a catch trial, a False Alarm (FA) was recorded and the rat received a 4-s timeout during which the house light was turned off and no trial could be initiated. Testing was carried out in a sound-attenuating chamber. Stimuli were calibrated using a sound level meter (Larson-Davis System 824) equipped with a half-inch microphone (Larson-Davis model 2520) at a location where the animal’s head would be during a trial.

**Figure 1:**
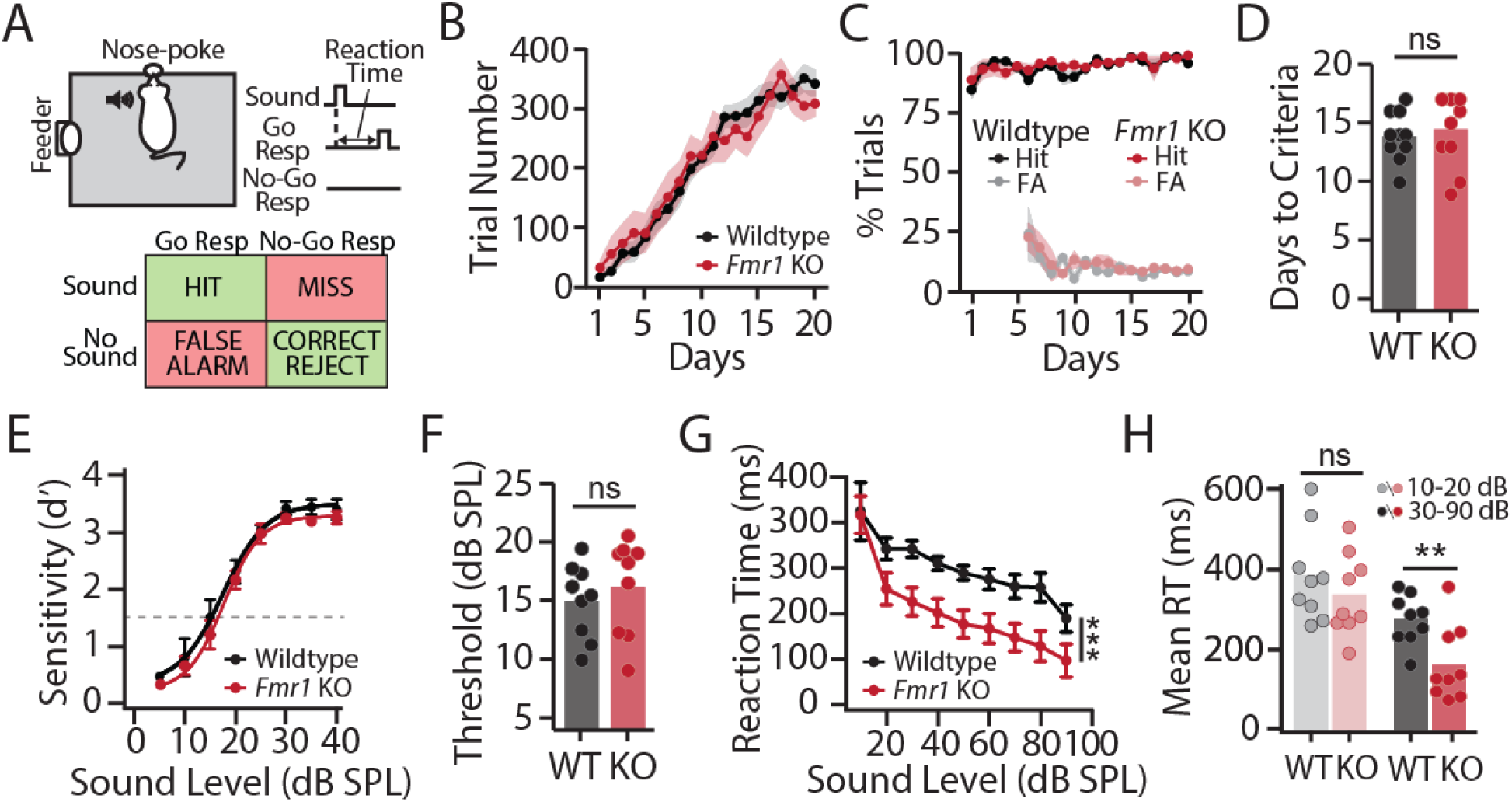
Psychoacoustic assessment of sound detection thresholds and suprathreshold auditory reaction times in male *Fmr1* KO and WT littermate rats. **(A)** Schematic of operant sound detection apparatus and paradigm. Food-restricted rats were trained to detect broadband noise (BBN) bursts using a Go/No-go paradigm. **(B)** *Fmr1* KO rats (red, n = 9) and littermate WT rats (black, n = 9) initiated the same number of trials over training days. **(C)** WT and KO rats generated the same percentage of HITs and FAs over training days. **(D)** WT and KO rats learned the task to the criteria (>200 trials/session, >90% correct, <15% FA, 5-consecutive days) over the same number of days. **(E)** Psychometric functions showing mean d’-prime values versus intensity for trained *Fmr1* KO and WT rats in response to broadband noise bursts (BBN; 1-42 kHz, 50 ms, 0-40 dB SPL, 5 dB steps). **(F)** No differences in hearing thresholds between KO and WT rats estimated using a criterion of d’ = 1.5 (dashed line in E). **(G)** Reaction time (RT)-intensity functions collected for BBN bursts (1-42 kHz, 50 ms, 10-90 dB SPL, 10 dB steps). RTs were significantly faster in *Fmr1* KO animals (\*\*\**p* < 0.0001). **(H)** Mean RTs at two lowest intensities (10-20 dB SPL) and five highest intensities (30-90 dB SPL) in WT and *Fmr1* KO rats. RTs in WT and KO rats are similar at low intensities, but significantly faster in KO rats at high intensities (\*\**p* < 0.001). For this and subsequent figures, data is plotted as mean +/− SEM. Bar graphs represent mean data with overlaid scatter plots of data from each individual animal.

Rats were initially trained to detect 60 dB SPL broadband noise (BBN, 1-42 kHz) bursts. Criteria for training was > 200 trials initiated, > 90% hit rate, and <15% false alarm rate over 5 consecutive days. Following training, sound intensity was varied using the Method of Constant Stimuli (MOCS); within each 10-trial block, seven target intensities were presented randomly along with three catch trials. To measure thresholds, noise bursts and tone bursts were presented at intensities from −5 to 45 dB SPL in 5 dB steps. Mean HIT and FA rates were used to calculate the sensitivity index d’ for each intensity and noise or tone burst detection thresholds were estimated using a conservative d’ value of 1.5 (Radziwon et al., 2019; Radziwon et al., 2009; Steckler, 2001). RT-I functions were collected for noise and tone bursts presented at intensities from 10-90 or 30-90 dB SPL in 10 dB steps. RT, defined as the time from sound stimulus onset to the time the rat removed its nose from the nose-poke hole, was assessed only for correct HIT trials (Fig 1A). To test for temporal integration of loudness, thresholds and RT-I functions were evaluated for BBN bursts of 50, 100, and 300 ms duration. To test for spectral integration of loudness, thresholds and RT-I functions were evaluated for 16 kHz tone and narrow band noise (NBN) bursts (300 ms) with nominal bandwidths of 1/3 octave (14.1 – 17.8 kHz), 1 octave (11.3 – 22.8 kHz), or 2 octaves (8 – 32 kHz). To test for frequency effects, thresholds and RT-I functions were evaluated for tone bursts (300 ms) at 4, 8, 16, 32 kHz. At least 600 trials (200 trials on 3 consecutive days of testing) were used to estimate each quiet threshold and loudness growth RT-I function in KO and WT rats.

### 2.4 MTEP Treatment

MTEP (Cayman Chemical, #14961), an mGlu5 receptor negative allosteric modulator, was dissolved in saline at 10 mg/ml. MTEP was administered intraperitoneal (i.p.) at 1 mg/kg, 3 mg/kg or 10 mg/kg, doses shown to be behaviorally effective while maintaining receptor specificity (Pilc et al., 2002; Spooren et al., 2000). Baseline RT-I functions for BBN (50 ms) were measured for 1-week in *Fmr1* KO and WT rats and then MTEP (1, 3 or 10 mg/kg) or saline was administered acutely 30 minutes before behavioral testing. Animals received each dose of MTEP or saline in pseudorandomized order while allowing for >1-week washout between treatments. Animals continued to be tested daily for RT-I functions between treatments. The selection of MTEP doses and timing of injections were based on the known half-life of MTEP in the brain (Anderson et al., 2002) and previous studies demonstrating this dosing regimen to be behaviorally effective (Spooren et al., 2000; Varty et al., 2005).

## 3 Results

### 3.1 *Fmr1* KO and WT rats exhibit similar learning and performance on an operant sound detection task

Male *Fmr1* KO (n=9) and WT (n=9) rats were trained to detect broadband noise bursts (BBN, 1-42 kHz, 50 ms, 5 ms rise/fall) using a Go/No-go operant conditioning paradigm (Fig. 1A). Correct Go responses were recorded as a HIT, failure to respond on Go trials were counted as a MISS, correct No-go responses were considered a Correct Rejection (CR), and incorrect Go responses on catch trials were recorded as a False Alarm (FA) (Fig 1A). During training, the mean (+/−SEM) number of trials increased over the first 15 training days and then plateaued in both genotypes and there was no significant difference between *Fmr1* KO and WT rats in terms of the number of trials initiated (Fig 1B). Two-way repeated measure ANOVA found a significant effect of training days on number of trials initiated (F_19, 304_ = 87.14, \*\*\**p* < 0.0001) but no significant effect of genotype (F_1, 304_ < 0.0001, *p* = 0.9994). There was also no significant difference between WT and KO rats in terms of percent of trials scored as HIT or percent scored as FA trials (Fig 1C). Two-way repeated measure ANOVA found a significant effect of training days on percent HIT (F_19,304_ = 3.274, \*\*\**p* < 0.0001) and percent FA (F_14, 120_ = 3.070, \*\**p* = 0.0012) but no significant effect of genotype on percent HIT (F_1, 304_ = 1.607, *p* = 0.2058) or percent FA (F_1, 120_ = 1.540, *p* = 0.2170). *Fmr1* KO and WT rats did not differ significantly on the number of days to reach the training criteria either (WT: 13.67 +/− 0.69 days; KO: 14.11 +/− 1.23; two-tailed t-test, t_16_ = 0.3614, *p* = 0.7225) (Fig 1D). Thus, WT and *Fmr1* KO rats learned the Go/No-go operant task at the same rate and to the same criteria. While FX individuals can often display differences in learning, motivation and impulsivity (Chromik et al., 2019; Schmitt et al., 2019), these results suggest that there are no genotype differences in these non-auditory factors in this sound detection task.

### 3.2 *Fmr1* KO rats have normal sound detection thresholds

To determine sound detection sensitivity in *Fmr1* KO and WT rats, BBN bursts (50 ms) were presented at near threshold intensities (5-45 dB, 5 dB steps) using the method of constant stimuli (MOC) (Radziwon et al., 2009). HIT and FA rates from over 600 trials (200 trials/day for 3 consecutive days) were used to determine d-prime (d’), a standard metric of sensitivity from signal detection theory (Steckler, 2001). Psychometric curves for *Fmr1* KO (n=9) and WT (n=9) rats were constructed by plotting d’ (mean+/−SEM) as a function of sound intensity (Fig 1E). d’ increased with sound intensity in both genotypes, indicating that as sounds grew louder they were more readily detectable, and psychometric functions from *Fmr1* KO and WT rats largely overlapped with no genotype differences in d’ across intensities (Fig. 1E). Two-way repeated measure ANOVA found a significant effect of intensity on d’ (F_7, 105_ = 52.64, \*\*\**p* < 0.0001), but no significant effect of genotype (F_1, 105_ = 0.5783, *p* = 0.4487). There was no significant difference in BBN thresholds between *Fmr1* KO and WT rat using a conservative threshold criterion of d’ = 1.5 (WT: 14.96 +/− 1.062 dB SPL; KO: 16.21 +/− 1.364 dB SPL; two-tailed t-test, t_16_ = 0.7234, *p* = 0.4799). These results indicate that *Fmr1* KO rats have comparable performance on a sound detection task to WT animals and normal hearing thresholds.

### 3.3 *Fmr1* KO rats exhibit excessive loudness growth

d’ is a sensitive measure of near threshold sound detection but saturates rapidly above threshold (Fig 1E). This metric therefore does not adequately convey suprathreshold loudness perception, which may be most affected in individuals with FX and ASD (Danesh et al., 2015; Gomes et al., 2008; Williams et al., 2021b). Because RT-I functions provide a valid measure loudness growth in humans and animals (Lauer and Dooling, 2007; Marshall and Brandt, 1980; May et al., 2009), auditory RTs were measured in response to BBN bursts (50 ms) from near threshold to suprathreshold sound intensities (10-90 dB SPL, 10 dB steps) in the same group of animals to determine if *Fmr1* KO rats showed signs of exaggerated loudness growth. Mean (+/−SEM) RTs values at 10 dB SPL, near the threshold of detectability, were approximately 400 ms for both Fmr1 KO (n=9) and WT (n=9) rats (Fig 1G). RTs became progressively faster with increasing intensity in both genotypes; however, the decrease in RT was much greater for *Fmr1* KO rats (Fig 1G) and RTs were significantly shorter in *Fmr1* KO rats compared to WT rats (Fig 1H). Two-way repeated measure ANOVA found a significant effect of intensity on RTs (F_8,144_ = 12.201, \*\*\**p* < 0.0001) and a significant effect of genotype (F_1, 144_ = 40.968, *\*\*\*p* < 0.0001), but there was no significant interaction effect (F_8,144_ = 0.607, *p* = 0.771). To visualize the RT differences between WT and KO at low versus high intensities, mean RTs were computed from 10-20 dB SPL and 30-90 dB SPL for the two genotypes (Fig. 1H). The mean RTs of WT and *Fmr1* KO rats at near threshold intensities, 10-20 dB SPL, were not significantly different (WT: 384.0 +/− 38.63 ms; KO: 335.8 +/− 33.72 ms; two-tailed t-test, t_16_ = 0.9407, *p* = 0.3609). However, at intensities from 30-90 dB SPL, the mean RTs of *Fmr1* KO rats were significantly faster than WT rats (WT: 275.0 +/− 20.51 ms; KO: 163.8 +/− 31.30 ms; two-tailed test, t_36_ = 2.971, \*\**p* = 0.009). Taken together, these results indicate that RT grows more rapidly at suprathreshold intensities in *Fmr1* KO rats than WT rats, suggestive of increased loudness growth.

### 3.4 Disrupted temporal Integration of loudness in *Fmr1* KO rats

Loudness perception not only depends on sound intensity but also duration, with the perceived loudness of a sound increasing with stimulus duration out to approximately 300 ms after which it remains constant (Buus et al., 1997; Florentine et al., 1998; Pedersen and Poulsen, 1973; Radziwon and Salvi, 2020). To determine if temporal integration of loudness was disrupted in *Fmr1* KO rats, RT-I functions were measured using BBN bursts of 50, 100 and 300 ms duration. In WT rats (n=9), mean (+/−SEM) RTs became significantly faster with increasing duration but maintained similar intensity-dependent changes, leading to RT-I functions that were roughly parallel but stacked above one another (Fig. 2A). RTs were fastest for 300 ms BBN bursts, slowest for 50 ms bursts, and intermediate for 100 ms bursts. Two-way repeated measure ANOVA found that RTs in WT rats became significantly faster with both intensity (F_6,112_ =3.91, *\*\*p =* 0.003) and duration (F_2, 112_ =80.55, \*\*\**p* < 0.0001), consistent with previous reports of temporal integration of loudness in normal rats (Radziwon and Salvi, 2020). RTs in *Fmr1* KO rats also decreased as intensity increased (Fig. 2B); however, there was no effect of sound duration on RTs in KO animals as the RT-I functions obtained with 50, 100 and 300 ms BBN bursts overlapped one another (Fig. 2B). Two-way repeated measure ANOVA found a significant effect of intensity on RT in Fmr1 KO rats (F_6, 112_ = 3.07, \**p* < 0.011) but not duration (F_2, 112_ = 2.95, *p* = 0.057). Importantly, there was little evidence of temporal integration of loudness in *Fmr1* KO rats even at low intensities where there is ample room for RTs to become faster with increasing duration. The lack of temporal integration is not due to a “floor” effect because RTs clearly became faster in *Fmr1* KO rats at higher sound intensities.

**Figure 2:**
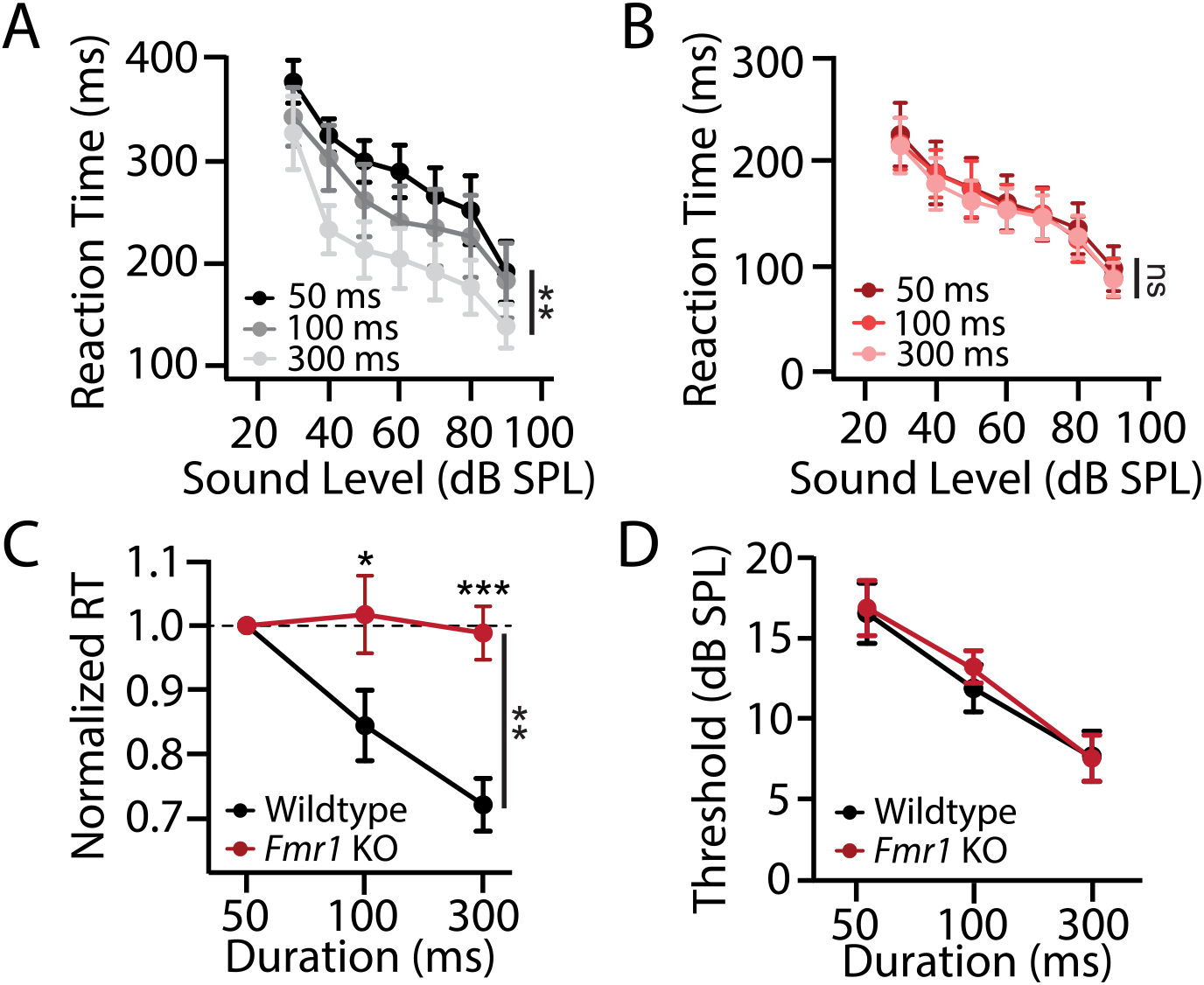
Disrupted temporal integration of loudness in *Fmr1* KO rats. Auditory reaction time-intensity (RT-I) functions measured with broadband noise bursts (BBN) of 50, 100 and 300 ms duration in **(A)** WT (+/− SEM, n=9) rats and **(B)** littermate *Fmr1* KO (+/−SEM, n = 9) rats. In WT rats, RTs decreased significantly with intensity (\*\**p*<0.0025) and duration (\*\*\**p <* 0.001) whereas in *Fmr1* KO rats, RTs only decreased significantly with intensity (\**p* < 0.0114), but not duration (*p* = 0.057). **(C)** Mean RTs across intensities (30-90 dB SPL) in WT and KO rats as a function of BBN duration, normalized to mean RTs for 50 ms BBN. RTs in WT rats were modulated by duration to significantly greater extent than *Fmr1* KO rats at 100 ms (\*\**p* < 0.01) and 300 ms (\*\*\**p* < 0.001). **(D)** Threshold of audibility (defined at d’ = 1.5) plotted as function of BBN duration. Thresholds decreased with duration in both *Fmr1* KO and WT rats and there was no significant difference in BBN thresholds across genotypes at any duration.

To quantify the relative effect of sound duration on RT across genotypes, normalized RTs were computed by dividing the average RT across intensities (30-90 dB) measured with 50, 100 ms and 300 ms BBN bursts by the average RT across intensities (30-90 dB) obtained with 50 ms BBN bursts for each rat. In the WT animals, mean (+/−SEM) normalized RT declined from 1.0 at 50 ms to approximately 0.7 at 300 ms, indicating that RTs at 300 ms were approximately 30% faster than at 50 ms (Fig. 2C). In *Fmr1* KO rats, mean normalized RTs were essentially unchanged from 50 ms (1.0) to 300 ms (0.99). Two-way ANOVA found a significant effect of duration (F_2, 32_ = 9.765, \*\*\**p* < 0.0001), a significant effect of genotype (F_1, 32_ = 11.27, \*\**p* = 0.0037), and a significant interaction between genotype and duration (F_2, 32_ = 8.503, \*\**p* = 0.001) on normalized RTs. Bonferroni post-hoc analysis found that duration has significantly more impact on average RT in WT animals compared to *Fmr1* KO rats at 100 ms (\**p* < 0.05) and 300 ms (\*\*\**p* < 0.0001). Thus, the lack of temporal integration of loudness in *Fmr1* KO rats indicates that faster RTs in *Fmr1* KO likely reflects a genuine perceptual disruption rather than effects due to non-auditory factors such as motivation or motor differences.

Temporal integration also affects the threshold of audibility. Hearing thresholds typically decrease 8-15 dB as stimulus duration increases out to approximately 300 ms, after which it remains constant (Pedersen and Salomon, 1977). To test for genotype differences in temporal integration at the threshold of audibility, sound detection thresholds were measured in WT and *Fmr1* KO rats using 50, 100 and 300 ms BBN bursts from 0-40 dB SPL. Mean (+/− SEM) thresholds in KO rats (n=9) were similar to those of WT rats (n=8) at 50, 100 and 300 ms (Fig. 2D). As duration increased from 50 to 300 ms, thresholds decreased from 16.9 +/− 1.87 dB SPL to 8.05 +/− 1.56 dB SPL in WT rats and 17.24 +/− 1.71 dB SPL to 7.93 +/− 1.42 dB SPL in KO rats. Two-way repeated measure ANOVA found that duration has a significant effect on thresholds (F _2, 30_ = 47, \*\*\**p*<0.0001); however, there was no significant effect of genotype (F_1, 30_ = 0.077, *p* = 0.785) and no significant interaction between genotype and duration (F_2, 30_ = 0.31, *p* = 0.735).

### 3.6 Enhanced spectral integration of loudness in KO rats

Loudness not only varies with intensity and duration, but also stimulus bandwidth (Cacace and Margolis, 1985; Scharf and Meiselman, 1977; Yost and Shofner, 2009; Zwicker et al., 1957). Loudness remains constant when the total energy of the stimulus lies within the critical band, but loudness increases as energy spreads outside the critical band. To test for spectral integration of loudness, RT-I functions were measured with 16 kHz tone bursts and 1/3, 1 and 2 octave-wide NBN bursts (300 ms) centered around 16 kHz. Mean (+/−SEM) RTs of WT rats (n = 8) decreased with intensity; however, the characteristics of the RT-I function were bandwidth dependent (Fig. 3A). The RT-I functions for 16 kHz and the 1/3 octave NBN were nearly identical indicating that these two stimuli were perceived as equally loud. However, as bandwidth increased further RTs became faster, indicative of spectral integration of loudness. Two-way repeated measure ANOVA revealed a significant effect of sound intensity (F_4, 84_ = 34.114, ***p < 0.0001) and bandwidth (F_3, 84_ = 4.324, \**p* = 0.016) on RT in WT animals. There was also a significant interaction between bandwidth and intensity (F_12, 84_ = 2.345, *\*p* = 0.012). To elucidate the interaction of bandwidth and intensity in WT rats, the mean RTs were plotted as a function of bandwidth at each intensity (Fig. 3B). RTs for 16 kHz tones (a nominal bandwidth of 1 Hz), 1/3 octave, and 1 octave wide NBN were not significantly different from one another. The only significant differences in RTs occurred at 30 and 40 dB SPL for the largest bandwidth separation (1 Hz vs 2 octave) (Bonferroni post-hoc, \**p*<0.05).

**Figure 3:**
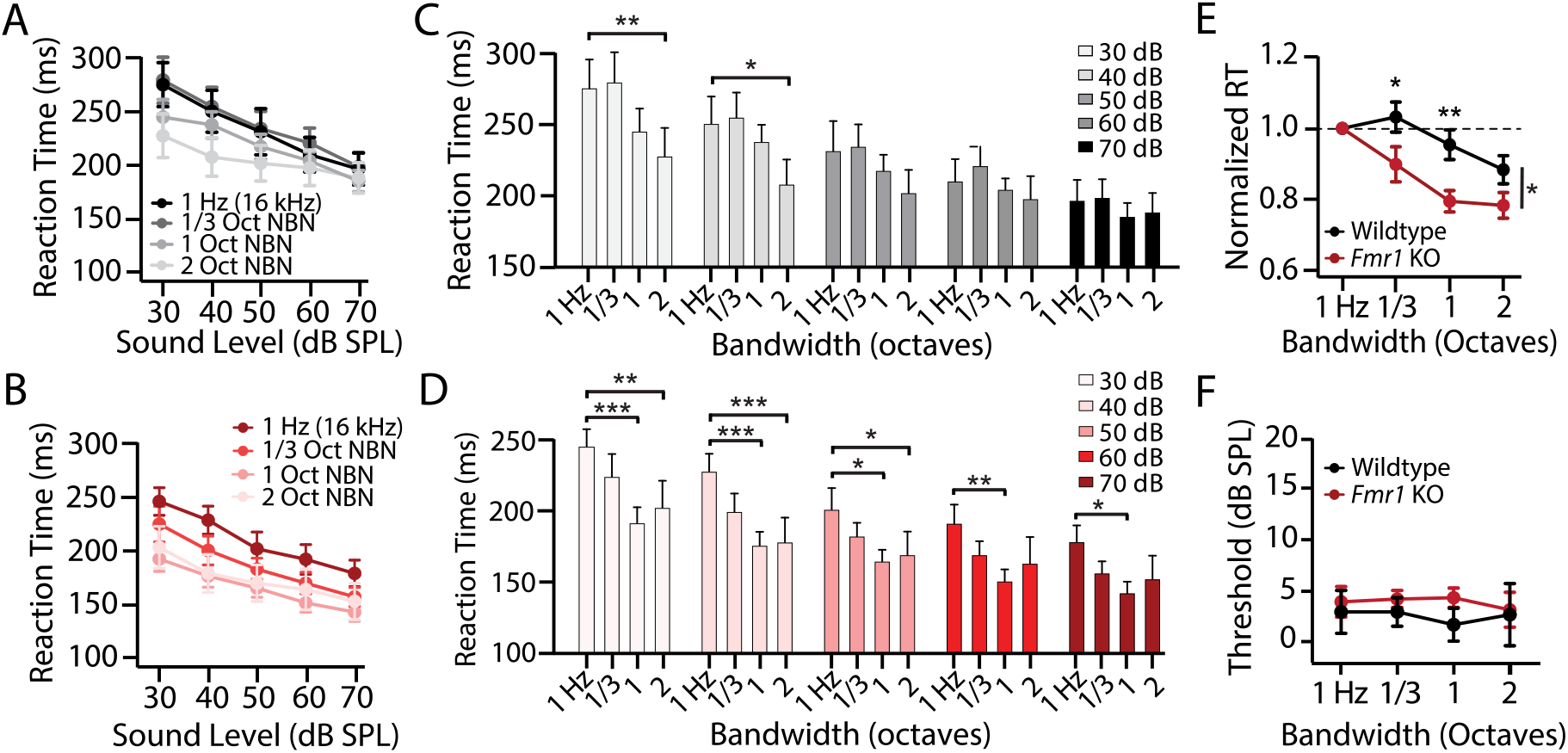
Altered spectral integration of loudness in *Fmr1* KO rats. **(A-B)** Mean (+/−SEM) reaction time-intensity (RT-I) functions in WT (n=8) and littermate *Fmr1* KO (n = 8) rats. Stimuli were 300 ms tone bursts (16 kHz, 1 Hz nominal bandwidth) and 300 ms narrow band noise (NBN) bursts centered at 16 kHz with bandwidths of 1/3, 1, or 2 octaves (Oct) from 30 to 70 dB SPL in 10 dB steps. **(A)** Mean (+/−SEM) RT-I functions in WT animals. RTs decreased significantly with intensity (***p < 0.0001) and bandwidth (\**p* = 0.016) and there was a significant bandwidth-intensity interaction (\**p*=0.012). **(B)** Mean (+/−SEM) RT-I functions in *Fmr1* KO animals. RTs decreased significantly with intensity (\*\*\**p* < 0.0001) and bandwidth (\*\**p* = 0.004) and the interaction between bandwidth and intensity not significant. **(C)** Mean RTs (+SEM) at each bandwidth (1 Hz, 1/3, 1, and 2 Oct) and each intensity (30-70 dB SPL) in WT animals. RTs at 1 Hz (16 kHz) were significantly slower from 2 Oct NBN at 30 and 40 dB SPL only (*p < 0.05). **(D)** Mean (+SEM) RTs at each bandwidth (1 Hz, 1/3, 1, and 2 Oct) and each intensity (30-70 dB SPL) in *Fmr1* KO animals. Significant differences in RT between 1 Hz (16 kHz) and 1 and/or 2 Oct NBN at all intensities (\**p* < 0.05, \*\**p* < 0.01, \*\*\**p* < 0.001). **(E)** Mean RTs from 30-70 dB SPL in WT and KO rats as a function of bandwidth, normalized to 1 Hz. The effect of bandwidth on RTs was significantly greater for *Fmr1* KO verse WT animals for 1/3 (*p < 0.05) and 1 (**p < 0.01) octave band noise. **(F)** Mean (+/−SEM) thresholds as a function of bandwidth in *Fmr1* KO and WT rats. Thresholds (defined at d’ = 1.5) were not affected by bandwidth and there was no significant different in thresholds across genotypes at any bandwidth.

In *Fmr1* KO rats (n=8), the mean (+/−SEM) RTs also became faster with increasing intensity and bandwidth; however, the effects of bandwidth on RT-I functions in KO animals were distinct from WT controls (Fig. 3C). While RT-I functions for two the two largest bandwidths (1 and 2 octaves) largely overlapped in KO animals, the functions were shifted upward from 1 Hz and 1/3 octave wide NBN (Fig. 3C). Two-way repeated measure ANOVA found a significant effect of sound intensity (F_4, 84_ = 68.174, \*\*\**p* < 0.0001) and bandwidth (F _3, 84_ = 6.155, \*\**p* = 0.004) in *Fmr1* KO rats, but there was no significant interaction between bandwidth and intensity (F_12, 84_ = 1.767, *p*=0.067). To visualize the effect of bandwidth, mean RTs were plotted as function of bandwidth from 30 to 70 dB SPL (Fig. 3C-D). At all intensities, there was a significant decrease in RT between 1 Hz (16 kHz tone) and 1/3 and/or 1.0 octave wide NBN (Bonferroni post-hoc, \**p* < 0.05).

To quantify the relative effect of bandwidth across genotypes, the mean RT (30-70 dB SPL) at each bandwidth was normalized to the mean RT at 1 Hz (i.e., 16 kHz). Mean normalized RTs are plotted as function of bandwidth for WT and *Fmr1* KO rats in Fig. 3E. While RTs generally became faster with increasing bandwidth in both genotypes, RTs in *Fmr1* KO animals were more sensitive to smaller bandwidth changes. Two-way repeated measures ANOVA found a significant effect of bandwidth (F_3, 42_ = 14.24, \*\*\**p* < 0.0001) and genotype (F_1, 42_ = 8.222, \**p* = 0.0124) on normalized RT and there was a significant interaction between bandwidth and genotype (F_3, 42_ = 2.879, \**p* = 0.047). Post-hoc analysis demonstrated that RTs were significantly faster in *Fmr1* KO animals at 1/3 (Bonferroni post-hoc, *p < 0.05) and 1 (Bonferroni post-hoc, **p < 0.01) octave NBN compared to WT littermates. These results show there is a much greater decrease in RT with increasing bandwidth in KO rats than WT rats and the decline occurs at narrower bandwidth, evidence of greater spectral integration of loudness in *Fmr1* KO rats.

### 3.7 No spectral integration at threshold

To test for genotype differences in spectral integration at the threshold of audibility, behavioral detection thresholds in quiet were determined for 300 ms tone burst at 16 kHz and the three NBN bandwidths. The mean thresholds (+/−SEM, n=8) in *Fmr1* KO and WT rats were not significantly different from one another (Fig. 3F). Two-way repeated measure ANOVA found no significant effect of genotype (F_1, 42_ = 0.441, *p* = 0.517) or bandwidths (F_3, 42_ = 0.144, *p* = 0.933) on detection thresholds and no interaction between the factors (F_3, 42_ = 0.299, p = 0.826). Thus, WT and *Fmr1* KO rats have similar thresholds and there was no difference in spectral integration at the threshold of audibility.

### 3.8 Loudness growth at different sound frequencies in WT and KO rats

To identify potential frequency-dependent differences in loudness growth between KO (n=9) and WT (n=8) rats, RT-I functions were evaluated at 4, 8, 16 and 32 kHz (Marshall and Brandt, 1980; Radziwon and Salvi, 2020). The mean (+/−SEM) RT-I functions assessed with 300 ms tone bursts decreased with intensity for both genotypes, but the RT-I functions for *Fmr1* KO rats were consistently below and roughly parallel to RT-I functions of WT rats at all frequencies (Fig. 4A-D). Two-way repeated measure ANOVAs found significant effects of intensity and genotype on RT-I functions at all frequencies, respectively (<u>4 kHz</u>: F_6, 105_ = 17.89, \*\*\**p* < 0.0001; F _1, 105_ = 22.21, *\*\*\*p* < 0.0001; <u>8 kHz</u>: F_6, 105_ =14.74, \*\*\**p* < 0.001; F_1, 105_ = 23.84, \*\*\**p* < 0.0001; <u>16 kHz</u>: F_6, 105_ =7.68, \*\*\**p* < 0.0001; F_1, 105_ = 10.92, \*\**p =* 0.0013; <u>32 kHz</u>: F_6, 105_ = 4.52, ** *p* = 0.0004; F_1,105_ = 14.47, \*\**p* = 0.0002). There were no significant interactions between intensity and genotype at any frequency.

**Figure 4:**
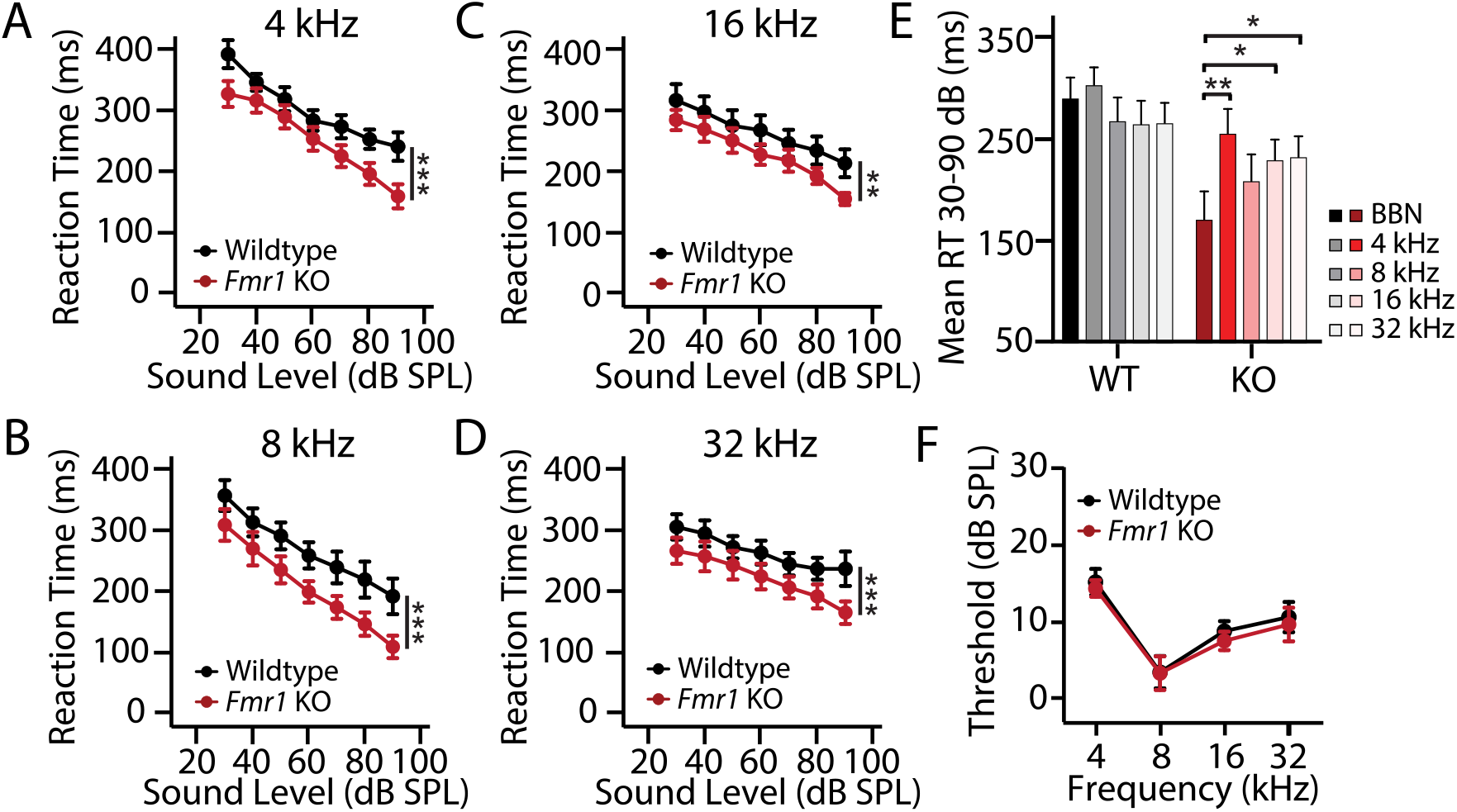
Loudness growth at different frequencies in WT and *Fmr1* KO rats. **(A-D)** Mean (+/−SEM) reaction time-intensity function in WT (n = 8) and KO (n = 9) rats obtained with 300 ms tone bursts at **(A)** 4 kHz, **(B)** 8 kHz, **(C)** 16 kHz and **(D)** 32 kHz. In both genotypes, RTs varied significantly as a function of sound intensity for all frequencies (\*\*\**p* < 0.0001). RTs were significantly faster in *Fmr1* KO rats than WT rats at 4 kHz (\*\*\**p* < 0.0001), 8 kHz (\*\*\**p* < 0.0001), 16 kHz (\*\**p* = 0.0013) and 32 kHz (\*\*\**p* = 0.0002). **(E)** Mean RTs across intensities (30-90 dB SPL) in response to BBN bursts (300 ms) and 4, 8, 16 and 32 kHz tone bursts (300 ms) in *Fmr1* KO and WT rats. Among WT rats, mean RT for BBN was not significantly different from mean RT for 4, 8, 16 or 32 kHz. Among *Fmr1* KO rats, mean RT for BBN was significantly faster than mean RTs at 4 (\*\**p* < 0.01), 16 (\**p* < 0.05) and 32 kHz (\**p* < 0.05). **(F)** Mean (+/−SEM) thresholds at 4, 8, 16 and 32 kHz for WT and *Fmr1* KO rats. Mean thresholds varied significantly with frequency (\*\*\**p* < 0.0001), but thresholds not significantly across genotypes at any frequency.

To determine if RTs to BBN bursts differed from tone bursts across genotype, mean RTs in WT rats (n=8) and KO rats (n=9) were computed from 30 to 90 dB SPL for BBN bursts (300 ms) and 4, 8, 16 and 32 kHz tone bursts (300 ms) (Fig. 4E). Mean RTs were significantly slower in WT rats than *Fmr1* KO rats across all stimulus conditions (BBN and 4 frequencies), as a two-way repeated measures ANOVA found a significant effect of genotype (F_1, 64_ = 5.54, *\*p* = 0.0318). However, the effect of stimulus condition (F_4, 64_ = 2.18, *p* = 0.081) and interaction of stimulus condition and genotype (F_4, 64_ = 1.97, *p* = 0.109) were not significant. Post-hoc analysis showed that the mean RTs for BBN bursts and 4, 8, 16 and 32 kHz tone bursts did not differ significantly from one another in WT rats. However, in *Fmr1* KO rats, the mean RT for BBN was significantly faster than the mean RTs at 4, 16 and 32 kHz tone bursts (Bonferroni post-hoc, \*\**p* < 0.01, \**p* < 0.05). These results suggest that loudness growth is altered across a broad range of sound frequencies in *Fmr1* KO rats and provide further evidence for greater spectral integration of loudness in these animals.

### 3.9 No differences in tone detection thresholds between *Fmr1* KO and WT rats

To determine if *Fmr1* KO and WT rats have similar pure tone sensitivity, thresholds were assessed at 4, 8, 16 and 32 kHz using tone bursts (300 ms). Mean (+/−SEM) thresholds for *Fmr1* KO (n=9) and WT (n=8) were lowest at 8 kHz and increased slightly at higher and lower frequencies. There was a significant main effect of frequency (F_3, 58_ =13.51, *p*<0.0001), but the thresholds for *Fmr1* KO and WT rats were not significantly different (F_1, 58_ =0.41, *p* = 0.527) and the interaction between genotype and frequency was not significant (F_3, 58_ = 0.04, *p* = 0.99). These results indicate that pure tone hearing thresholds are similar in *Fmr1* KO and WT rats.

### 3.10 mGlu5 inhibition selectively slows RTs in *Fmr1* KO but not WT rats

We next sought to determine if we could reverse auditory RT differences in *Fmr1* KO animals through pharmacological manipulation. Because dysregulated mGlu5 signaling is a core component of FX pathophysiology (Bhakar et al., 2012; Michalon et al., 2012; Pop et al., 2014), we determined if mGlu5 inhibition could restore normal loudness perception in *Fmr1* KO rats. Following baseline measurements, RT-I functions in response to 50 ms BBN bursts were measured in WT and *Fmr1* KO rats treated with the specific mGlu5 negative allosteric modulator MTEP at three doses (1, 3 and 10 mg/kg, i.p.) or saline 30 minutes prior to behavioral testing. RT-I functions in MTEP treated WT rats (+/−SEM, n=10) were nearly identical to those obtained at baseline or after control treatment with saline at all doses tested (Fig. 5A). Two-way repeated measure ANOVA did reveal a significant effect of treatment (F_4, 216_, \**p =* 0.025) in addition to intensity (F_6, 252_ = 26.243, \*\**p =* 0.001). However, post-hoc analysis determined that there was no significant difference between RT-I functions obtained following MTEP treatment at any dose compared to baseline or saline control conditions (Bonferroni post hoc, *p* > 0.05). There was a much clearer effect of MTEP treatment on *Fmr1* KO rats (n = 12), as MTEP dose-dependently upshifted RT-I functions in these animals. Two-way repeated measure ANOVA revealed a significant effect of intensity (F_6, 288_ = 62.114, \*\*\**p* < 0.0001) and treatment (F_4, 288_ = 13.399, \*\*\**p* < 0.0001). Moreover, post-hoc analysis revealed that RTs at nearly every intensity were significantly slower in *Fmr1* KO rats following the 3 or 10 mg/kg dose of MTEP relative to baseline and saline control conditions (Bonferroni post-hoc, \**p* < 0.05, \*\**p* < 0.001, \*\*\**p* < 0.0001).

**Figure 5:**
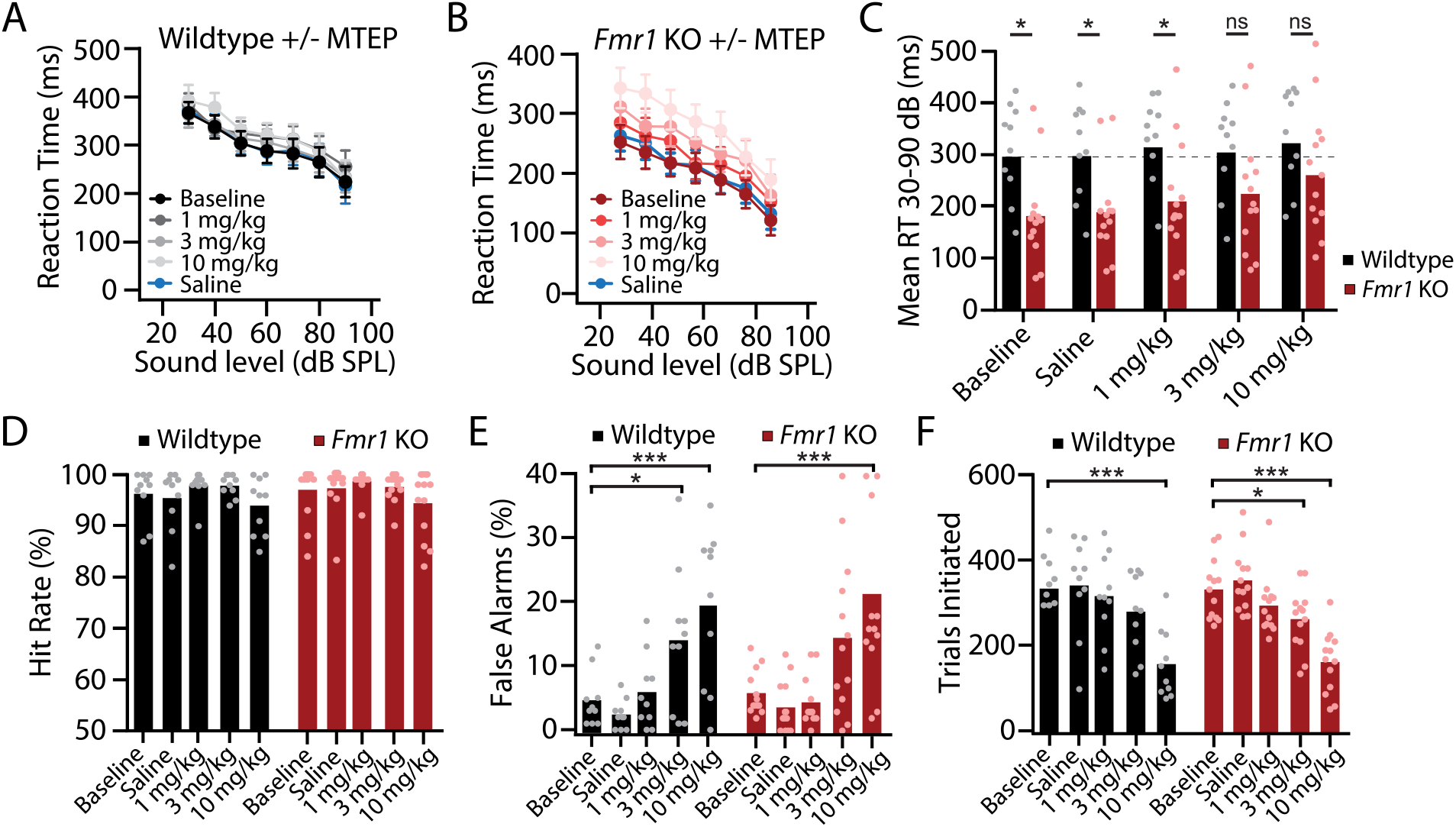
Inhibition of mGlu5 dose-dependently increases RTs in *Fmr1* KO but not WT rats. **(A)** Mean (+/−SEM, n = 10) reaction time-intensity (RT-I) functions from WT rats treated with different doses of the selective mGlu5 negative allosteric modulator MTEP or saline versus baseline control. Analysis of the RT-I functions revealed a significant effect of intensity (\*\*\**p* < 0.0001) and treatment (\**p* < 0.025); however, RTs obtained with MTEP treatments were not significantly different from baseline or saline control conditions. **(B)** Mean (+/−SEM, n = 12) RT-I functions from *Fmr1* KO rats treated with different doses of MTEP or saline versus baseline control. Analysis of the RT-I functions revealed a significant effect of intensity (\*\*\**p <* 0.0001) and treatment (\*\*\**p* < 0.0001) and RT-I functions were significantly different from baseline and saline control conditions when treated with 3 mg/kg (\*\**p* = 0.004) or 10 mg/kg (\*\*\**p* < 0.001) MTEP. **(C)** Mean RTs from 30 to 90 dB SPL in WT and *Fmr1* KO rats for baseline, saline, and 1, 3 and 10 mg/kg MTEP conditions. RTs were significantly faster in *Fmr1* KO rats compared WT rats for the baseline, saline control and 1 mg/kg MTEP conditions (\**p* < 0.05). Mean values for WT and *Fmr1* KO rats were not significantly different from one another for the 3 and 10 mg/kg MTEP treatments (*p* > 0.05). **(D)** Mean percent correct responses did not differ across experimental conditions in WT (*p* = 0.107) or *Fmr1* KO rats (*p* = 0.102). **(E)** Mean percent false alarms in WT rats and *Fmr1* KO rats varied significantly across treatment (\*\*\**p*<0.0001) but there was no significant effect of genotype (*p =* 0.703). False alarm rates were significantly greater with 3 mg/kg (\**p* < 0.05) and 10 mg/kg (\*\*\**p* < 0.001) MTEP treatment in WT animals and were significantly greater in KO animals at 10 mg/kg MTEP treatment (\*\*\**p* < 0.001) compared to baseline. **(F)** Mean trials initiated in WT rats and *Fmr1* KO rats varied significantly across treatment (\*\*\**p* < 0.0001) but there was no significant effect of genotype (*p =* 0.8496). Trial number was significantly decreased with 10 mg/kg (\*\*\**p* < 0.001) MTEP treatment in WT animals and was significantly decreased in KO animals with 3 mg/kg (\**p* < 0.05) and 10 mg/kg (\*\*\**p* < 0.001) MTEP treatment compared to baseline. Bar graphs represent mean data with overlaid scatter plots of data from each individual animal.

To assess genotype differences with MTEP treatment, average RTs were computed for intensities between 30 and 90 dB SPL and the mean values compared across genotype for baseline, saline, 1, 3 and 10 mg/kg MTEP (Fig. 5C). The average RTs in WT rats were relatively constant across treatments. In contrast, RTs in *Fmr1* KO rats dose-dependently increased with MTEP dose so that there was no significant difference in average RT between WT and KO rats treated with 3 or 10 mg/kg MTEP. Two-way repeated measure ANOVA revealed a significant effect of genotype (F_1, 84_ = 6.64, \**p* < 0.018), treatment (F_4, 84_ = 13.51, \*\*\**p* < 0.0001) and a significant genotype x treatment interaction (F_4, 84_ = 3.98, \*\**p* = 0.0052). The average RTs were significantly faster in *Fmr1* KO rats for the baseline, saline control, and 1 mg/kg MTEP conditions (Bonferroni post-hoc, \**p* < 0.05). However, the mean values for WT and *Fmr1* KO rats were not significantly different from one another for the 3 and 10 mg/kg MTEP treatments, suggestive of normal loudness restoration in *Fmr1* KO rats (Bonferroni post-hoc, *p* > 0.05).

To test for non-specific drug effects, mean percent HIT, mean percent FA and mean number of trials were examined for WT and *Fmr1* KO rats for all experimental conditions. Mean HIT rates in WT rats (Fig. 5D, left panel) and *Fmr1* KO rats (Fig. 5D, right panel) did not differ across experimental conditions in either WT (one-way repeated measure ANOVA; F_4, 36_ = 2.060, *p* = 0.1065) or *Fmr1* KO rats (one-way repeated measure ANOVA; F_4, 48_ = 2.050, *p* = 0.1022). However, FA rates dose-dependently increased in both WT (one-way repeated measures ANOVA; F_4, 36_ = 12.44, \*\*\**p* < 0.0001) and *Fmr1* KO (one-way repeated measures ANOVA; F _4, 36_ = 11.07, \*\*\**p* < 0.0001) (Fig 5E). In both genotypes, FA rate was significantly higher following 3 mg/kg and 10 mg/kg MTEP conditions than in the baseline and/or saline control conditions (Bonferroni post hoc \**p* < 0.05, \*\**p* < 0.001, \*\*\**p* < 0.0001). Mean number of trials per session also decreased at higher doses of MTEP in both WT (one-way repeated measures ANOVA; F _4, 36_ = 12.04, \*\*\**p* < 0.0001) and *Fmr1* KO rats (one-way repeated measures ANOVA; F _4, 48_ = 24.64, *\*\*\*p* < 0.0001) (Fig 5F). Both WT and *Fmr1* KO animals initiated significantly less trials following 10 mg/kg MTEP compared to baseline and saline control treatment (Bonferroni post hoc \**p* < 0.05, \*\**p* < 0.001, \*\*\**p* < 0.0001). While MTEP treatment affected trial initiation and FA rate equally in WT and KO animals, RTs were only affected in *Fmr1* KO rats, suggesting that the reversal of RT difference was not due to non-specific drug effects. While these results indicate that mGlu5 inhibition can reverse loudness disturbances in *Fmr1* KO rats, they also suggest that this treatment is associated with dose-limiting side effects that affected both genotypes equally.

## 4 Discussion

Here we used a perceptual decision-making task to perform a detailed characterization of sound intensity processing and loudness perception in a *Fmr1* KO rat model of FX. *Fmr1* KO animals learned the Go/No-go sound detection task at the same rate as WT counterparts and reached similar peak performance (Fig 1A-F). Despite similar sound detection levels, *Fmr1* KO rats responded with significantly faster RTs to stimuli across a wide-range of intensities, indicative of loudness hyperacusis (Fig 1G-H). To gain insights into the nature of this perceptual disturbance, we evaluated how auditory thresholds and suprathreshold RT-intensity functions were affected by sound duration, frequency and stimulus bandwidth in *Fmr1* KO and WT littermates. *Fmr1* KO animals exhibited impaired temporal integration of loudness when stimulus duration was increased (Fig 2), but enhanced spectral integration of loudness as bandwidth increased (Fig 3, 4). Importantly, we demonstrated that loudness hyperacusis in *Fmr1* KO rats could be normalized by mGlu5 antagonism, demonstrating that this auditory perceptual phenotype is related to a core molecular pathology of the disorder (Fig 5). However, the doses of MTEP that normalized loudness perception in *Fmr1* KO rats also had mild effects on task performance in both WT and KO rats, indicating the potential for dose-limited side effects of broad-spectrum mGlu5 inhibitors. Our results provide the first detailed behavioral characterization of a debilitating auditory phenotype (loudness hyperacusis) in an animal model of FX. These quantitative and clinically translatable behavioral assays provide researchers with a powerful experimental tool that can be used to identify effective therapies to treat one of the major sensory disabilities associated with FX and ASD.

### 4.1 Abnormal loudness perception in FX and ASD

Decreased sound tolerance is a common feature of FX and ASD, with prevalence rates between 75 and 85% (Williams et al., 2021b). The specific perceptual attributes underlying these sound tolerance disturbances remain unclear, as they could reflect disruptions to low-level sound processing, an altered ability to gate sensory input, and/or aberrant emotional responses to auditory stimuli (Williams et al., 2021a). While recent EEG studies in FX individuals and *Fmr1* KO mice have found neurophysiological evidence for low-level sound processing deficits, including increased magnitude of sound-evoked responses (Ethridge et al., 2016; Lovelace et al., 2018), auditory perceptual deficits in FX have not been well-characterized. Here, we found that auditory RTs decreased with intensity in both *Fmr1* KO and WT rats, but that RTs were consistently faster in *Fmr1* KO rats than WT littermates. The relationship between RT and intensity have been well documented in psychoacoustic studies (Lauer and Dooling, 2007; Marshall and Brandt, 1980; May et al., 2009) with RT-I functions being tightly correlated with loudness growth functions (Marshall and Brandt, 1980; Schlittenlacher et al., 2014; Wagner et al., 2004). These results therefore suggest that suprathreshold sounds are perceived as louder in *Fmr1* KO rats compared to WT littermates, indicative of loudness hyperacusis (Lauer and Dooling, 2007; Tyler et al., 2014). Because RTs in *Fmr1* KO and WT rats did not differ at low-intensities near threshold, faster RTs in *Fmr1* KO at suprathreshold intensities likely reflect aberrant loudness processing rather than motor differences. This interpretation is further supported by the fact that *Fmr1* KO rats exhibited altered temporal and spectral integration of sound intensity. If RT differences were due to non-auditory factors, then it would be expected that RTs in *Fmr1* animals would be modulated by changes to sound duration and bandwidth in a similar manner to their WT counterparts, but this was clearly not the case (Fig 2,3). Because the perceptual integration of sound duration and bandwidth were altered in opposite directions in *Fmr1* KO rats, this suggests that abnormal RTs in these animals are evidence of aberrant auditory processing rather than generalized hyperactivity or altered motor responses. These findings are consistent with previous studies showing lower loudness discomfort levels and steeper loudness growth functions among individuals with ASD (Demopoulos and Lewine, 2016; Khalfa et al., 2004; Rosenhall et al., 1999; Steigner and Ruhlin, 2014). Thus, sound tolerance issues in FX may be due in part to increased loudness perception.

### 4.2 Aberrant temporal and spectral integration of loudness

Loudness increases with stimulus duration out to ~300 ms (Zwislocki, 1969). In WT rats, temporal integration of loudness was expressed as a systematic decrease in RT from 50 to 300 ms (Fig 2A), consistent with previous results (Radziwon and Salvi, 2020). RTs in *Fmr1* KO rats, however, exhibited minimal RT changes as stimulus duration increased (Fig 2B). Thus, loudness processing is not only disrupted in the intensity domain, but also in the temporal domain. However, temporal integration was clearly present at low intensities near the threshold of audibility in *Fmr1* KO rats (Fig 2D), suggesting that the mechanisms responsible for temporal integration near threshold are different from those mediating temporal summation of loudness. Consistent with this notion, in humans with cochlear hearing loss, temporal summation of loudness remains relatively normal (Buus et al., 1999; Pedersen and Poulsen, 1973) whereas temporal integration at the threshold is significantly reduced (Hall and Fernandes, 1983; Plack and Skeels, 2007).

Loudness remains constant as long as the energy in the stimulus is within the critical band, but increases at wider bandwidths (Scharf, 1978; Zwicker et al., 1957). In WT rats, RTs became faster for bandwidths ≥ 1 octave suggesting a critical bandwidth between 1/3 to 1 octave (Fig. 3E). A significant decrease in RTs was already evident at 1/3 octave in *Fmr1* KO rats signifying that the critical bandwidth was ≤1/3 octave in these animals (Fig 3E). Because the critical band is narrower in *Fmr1* KO rats than WT rats, BBN would be perceived as louder in *Fmr1* rats. This would also explain why RTs are so much faster for BBN than for tone bursts in *Fmr1* rats (Fig 4E). The difference between *Fmr1* KO and WT rats for spectral integration of loudness is unlikely due to cochlear dysfunction because *Fmr1* KO rats have normal hearing thresholds and because cochlear hearing loss leads to a broadening of the critical band rather than a narrowing (Zwicker et al., 1957). Thus, enhanced spectral integration and diminished temporal summation of loudness in *Fmr1* rats are likely to be central rather than cochlear in origin, potentially due to imbalances in the magnitude, timing and spectral integration of excitatory and inhibitory inputs to central auditory neurons (Isaacson and Scanziani, 2011; Wehr and Zador, 2003). These rudimentary perceptual changes are not only clinically-relevant phenotypes that can be used to uncover potentially generalizable pathophysiological mechanisms and treatment strategies for FX, but they are also likely to be directly related to more complex phenotypes in FX and ASD, such as impaired language processing and communication. More detailed psychophysical studies should be conducted to determine if the novel disturbances in temporal and spectral integration observed in *Fmr1* KO rats occur in humans with FX and ASD.

### 4.3 Using RT-I functions as a drug-discovery and clinical outcome measure

FX has been a bellwether for illustrating the potential and pitfalls of translating pre-clinical findings into treatments for neurodevelopmental disorders (Berry-Kravis et al., 2017; Leigh et al., 2013; Nickols and Conn, 2014). Despite tremendous insight into the pathophysiological mechanisms of FX from animal models, clinical trials targeting identified molecular disturbances in FX have been disappointing to date (Berry-Kravis et al., 2018). For instance, the mGluR theory of FX has been the most influential model for understanding FX pathophysiology (Bear et al., 2004), positing that loss of FMRP leads to exaggerated protein synthesis linked of mGlu5 activation, resulting in altered synaptic function that is the root cause of cognitive impairment in FX (Bear et al., 2008; Bhakar et al., 2012). Decreasing mGlu5 activity has indeed been successful at reversing numerous phenotypes in animal models of FX (Dolen et al., 2007; Michalon et al., 2012). Despite this preclinical success, recent large-scale clinical trials targeting this receptor have largely failed (Berry-Kravis et al., 2016). Some of these clinical translation difficulties are due to the fact that broad-spectrum mGlu5 antagonists are associated with dose-limiting side effects and drug-dependent tolerance (Berry-Kravis et al., 2018; Erickson et al., 2017). A better understanding of how mGlu5 couples to FMRP-regulated protein synthesis might circumvent some of these issues by allowing for the development of pharmacotherapies that specially target signaling cascades thought to be involved in FX pathogenesis while leaving other side-effect producing signaling arms unaffected (McCamphill et al., 2020; Stoppel et al., 2017). Another key factor complicating the translation of apparently effective therapies in animal models into the clinic is that many animal phenotypes do not have equivalent correlates in humans, limiting their translational value (Berry-Kravis et al., 2018). The RT-I functions used in our study to assess loudness growth and hyperacusis have been carefully validated in human psychophysical studies (Lauer and Dooling, 2007; Marshall and Brandt, 1980). RT-I functions may thus provide researcher with a relatively simple, quantitative, and robust behavioral read-out for investigating auditory processing deficits in FX, which is a common and clinically important sensory phenotype that affects up to 85% of individuals with FX and ASD (Danesh et al., 2015; McCullagh et al., 2020; Williams et al., 2021b). Preclinical pharmacological studies in *Fmr1* KO rats can be used to carry out dose-response studies with prospective therapeutic compounds to test their effectiveness in reversing loudness disruptions and to identify non-specific side effects. Successful preclinical studies would be directly translatable to human studies aimed at suppressing loudness hyperacusis and possibly other sensory hypersensitivity disorders associated with FX and ASD, as well as other clinical disorders with high prevalence of hyperacusis such as Williams syndrome and fibromyalgia (Miani et al., 2001; Suhnan et al., 2017; Zarchi et al., 2015). To this end, we attempted to validate RT-I functions as a tool for screening drug therapies by determining the effect of mGlu5 inhibitors on RT differences in FX animals.

We found that MTEP, a selective mGluR5 negative allosteric modulator, dose-dependently normalized RT-I functions in *Fmr1* KO rats (Fig 5). Our results are therefore consistent with previous studies showing that mGlu5 inhibition reverses other auditory phenotypes, such as increased propensity for audiogenic seizures and altered acoustic startle response (Yan et al., 2005). However, the operant behavioral task used in this study also has several key features, such as self-initiated trials and no stimulus catch trials, that allow for monitoring of non-specific side effects of mGlu5 inhibition. Although MTEP normalized RT-I functions in *Fmr1* KO rats, it also affected other behavioral metrics in both WT and KO animals. These include a modest dose-dependent decrease in total trials initiated, suggestive of decreased motivation or appetite, and a dose-dependent increase in FA rates, suggestive of increased impulsivity or decreased attention. It is unlikely that MTEP-dependent changes to RT in FX animals were a byproduct of these changes in task performance, as RT was only affected in *Fmr1* KO rat but not WT littermates, whereas trial number and FA rate were equally affected in both genotypes (Fig 5). However, these side effects begin to appear at the same doses where the beneficial effects on RT are observed, indicating that broad spectrum mGluR5 antagonists may have a very narrow therapeutic range, limiting their clinical efficacy. These results demonstrate the utility of this task design in terms of screening pharmacotherapies for auditory phenotypes and identifying potential side-effects. Future studies can use this paradigm to optimize dosing structure, assess long-term tolerance development, and examine other potential therapies for FX and ASD. Importantly, we have shown that RT-I functions remain stable over several months (Radziwon and Salvi, 2020), suggesting this approach can be used for longitudinal studies.

## 4.5 Conclusion

Using RT-I functions to quantify loudness growth, we show for the first time that male *Fmr1* rats, in comparison to their WT littermates, show robust evidence of loudness hyperacusis to a variety of acoustic stimuli. Loudness hyperacusis in *Fmr1* KO rats was accompanied by enhanced spectral integration of loudness and deficient temporal summation of loudness as suprathreshold intensities. *Fmr1* KO rats had normal hearing thresholds and exhibited normal temporal integration at the threshold of audibility, perceptual characteristics compatible with normal cochlear function, but suggestive of central auditory processing deficits possibly mediated by an imbalance between excitation and inhibition. MTEP, an mGluR5 negative allosteric modulator, dose-dependently restored normal loudness growth in *Fmr1* KO rats but had no effect on RT-I measures of loudness growth in WT littermates. Behavioral RT-I measures of loudness growth thus represent a powerful tool for characterizing various dimensions of aberrant auditory processing in FX and ASD and can be used for preclinical screening of pharmacotherapies to treat loudness intolerance disorders and identifying potential side effects. These same psychophysical tests of loudness perception may also prove useful in the diagnosis or treatment of hyperacusis in FX and ASD individuals.

## List of abbreviations

FX: fragile X;
WT: wild type;
KO: knockout,
ASD: autism spectrum disorder;
RT-I: reaction time-intensity

## Acknowledgements

This research was supported in part by grants from NIH (RS: R21DC017813; BDA: F32DC015160, K01DC018310), The Simons Foundation for Autism Research Initiative (RS), and a NARSAD Young Investigator Award (BDA).

